# A Binary RNA and DNA Self-Amplifying Platform for Next Generation Vaccines and Therapeutics

**DOI:** 10.1101/2022.12.06.519322

**Authors:** Wilfred A. Jefferies, Kyung Bok Choi, Paolo Ribeca, Suresh Kari, Jay Young, Elizabeth Hui, Simon Yong Qi, Emmanuel Garrosvillas, Pamela Lincez, Tracy Welch, Iryna Saranchova, Cheryl G. Pfeifer

**Author notes:** Contributed equally (Paolo Ribeca, Suresh Kari, Jay Young).

## Abstract

Conventional mRNA-based vaccines were instrumental in lowering the burden of the pandemic on healthcare systems and in reducing mortality. However, such first-generation vaccines have significant weaknesses. Here, we describe a high-performance binary recombinant vectoral platform offering the flexibility to be used as a self-amplifying mRNA or a self-amplifying DNA. Both formats drive long-lasting expression and actuate robust antibody responses against SAR-CoV-2 spike, and neither format require encapsulation with lipid nanoparticles (LNP) in the generation immune responses. The platform combines the power of conventional mRNA with the low-dosage of self-amplifying vectors together with the simplicity, rapid creation, ease of storage, and convenience of distribution of plasmid DNA vectors. This platform promises to pave the way for more effective, less expensive, and truly democratized vaccines and therapeutics.

**One-Sentence Summary:** Gemini: a versatile platform that improves on existing vaccine formats in terms of effectiveness, manufacturing, distribution, and cost.

## MAIN TEXT

The zoonotic transmission of pathogens from animals to humans has become the kindling of emergent diseases, with the frequency dramatically increasing in our shift from hunters and gatherers to agrarian societies (1). Human interactions with animals through hunting, animal farming, trade of animal-based foods, wet markets or exotic pet trade (2), together with increased human interactions through global trade and travel (3) have ignited the fires of global pandemics. In the 20^th^ century alone, five major pandemics emerged, including, Smallpox, HIV/AIDS (1976), the sixth cholera pandemic (1899, 1923), the Spanish flu (1918 to 1920), and the Swine flu (2009), that resulted in over 100 Million deaths world-wide (4). Yet in the 21^st^ century, despite the sparks created by SARS-CoV-1 (2003), the global zeitgeist remained in the dark and was unprepared for the bonfire that became SARS-CoV-2 (2019). If not for the promethean intervention of the biotechnology community, combined with truly herculean efforts of public health authorities to collectively quell the flame of the COVID-19 pandemic through the swift introduction of first-generation SARS-CoV-2 vaccines, humanity would likely have been reduced to ashes, and may yet become a mere ember, in the absence of better vaccines (5). The first-generation vaccines were developed under an unprecedented, accelerated scheme underpinned by breakneck production and delivery rollout, with only a few months elapsing from design to testing and approval, and by in large, target the spike protein of SARS-CoV-2. The initial SARS-CoV-2 vaccines to appear on the world stage mainly differ in their underlying delivery platforms: Oxford/Astra Zeneca vaccine is based on a recombinant adenovirus vector, while Moderna and Pfizer vaccines are both based on a mRNA-based platform. Additional vaccine designs based on various formulation principles have since been developed in a number of countries, which include Novavax, a protein-based design containing spike and Valneva, a vaccine containing the inactivated virus (6). mRNA delivery systems have offered the advantage of rapid development of vaccines (7-9). This platform has been shown to be safe, effective, and adaptable for a variety of therapeutic applications (7-9). However, mRNA systems have been limited by their requirement for highly technical manufacturing, their inherent thermal instability (10) and their inefficient *in vivo* delivery in the absence of lipid nanoparticles (LNPs) (11). Overall, there remains a societal need to create new and more effective platforms with the clear aim of achieving sterilizing immunity.

Self-amplifying RNA (saRNA) delivery platforms stand out as leading technologies in vaccine development with the potential to solve many of the issues that have been highlighted for other platforms. Recombinant saRNA expression vectors featured an engineered replicon that can encode and drive high levels of antigen expression (12). Very low doses (micrograms) compared to mRNA technologies are required as tens of thousands of copies of saRNAs are synthesized directing immense amounts of payload mRNA transcription within recipient cells (12), and furthermore, saRNA vaccines can be delivered relatively noninvasively by intramuscular injection, similar to mRNA or DNA vaccines (12). Self-amplifying vaccines are considered safe and capable of inducing humoral and cellular immunity and they can also avoid the induction of anti-vector immunity, while simultaneously reducing the risk of the vector genome integration into the host genome (12). Manufacturing advantages when compared with conventional vaccines include a lower intrinsic risk of contamination with live infectious reagents and a much better scalability when mass production is required. On the other hand, the current generation of saRNA vectors also shares a number of issues with conventional mRNA platforms: they require technically demanding production processes involving *in vitro* transcription; they are generally unstable during long-term storage, and conventional dogma suggests they rely upon costly and technically demanding LNP encapsulation to allow uptake into cells where they express their payload proteins (12).

Here we report the creation and testing of Gemini, a bi-functional replicon platform that can be used in either a self-amplifying RNA (Gemini-R) or in a self-amplifying DNA (Gemini-D) plasmid (saDNA) format. Gemini addresses many of the problems with recent and traditional vaccine design and has perspicuous applications to SARS CoV-2 vaccines and broader applications to deliver and express other recombinant proteins or RNAs.

## Results

### Both Gemini-R and Gemini-D induce amplification in transfected cells

Gemini is a replicon-based self-amplifying dual expression vector containing a prokaryotic T7 promoter that can drive the transcription of mRNAs *in vitro* (Gemini-R) for uses in cells or tissues combined with a eukaryotic promoter that can faithfully transcribe mRNA after being delivered as plasmid DNA (Gemini-D) in cells or tissues (Figure 1). As an initial validation of the fidelity of the Gemini design, we confirmed active amplification by using RT-PCR to measure the expression of negative-stranded mRNA in HEK293 cells transfected with either Gemini-R or Gemini-D (Figure 2A, see also Materials and Methods).

**Fig 1.**
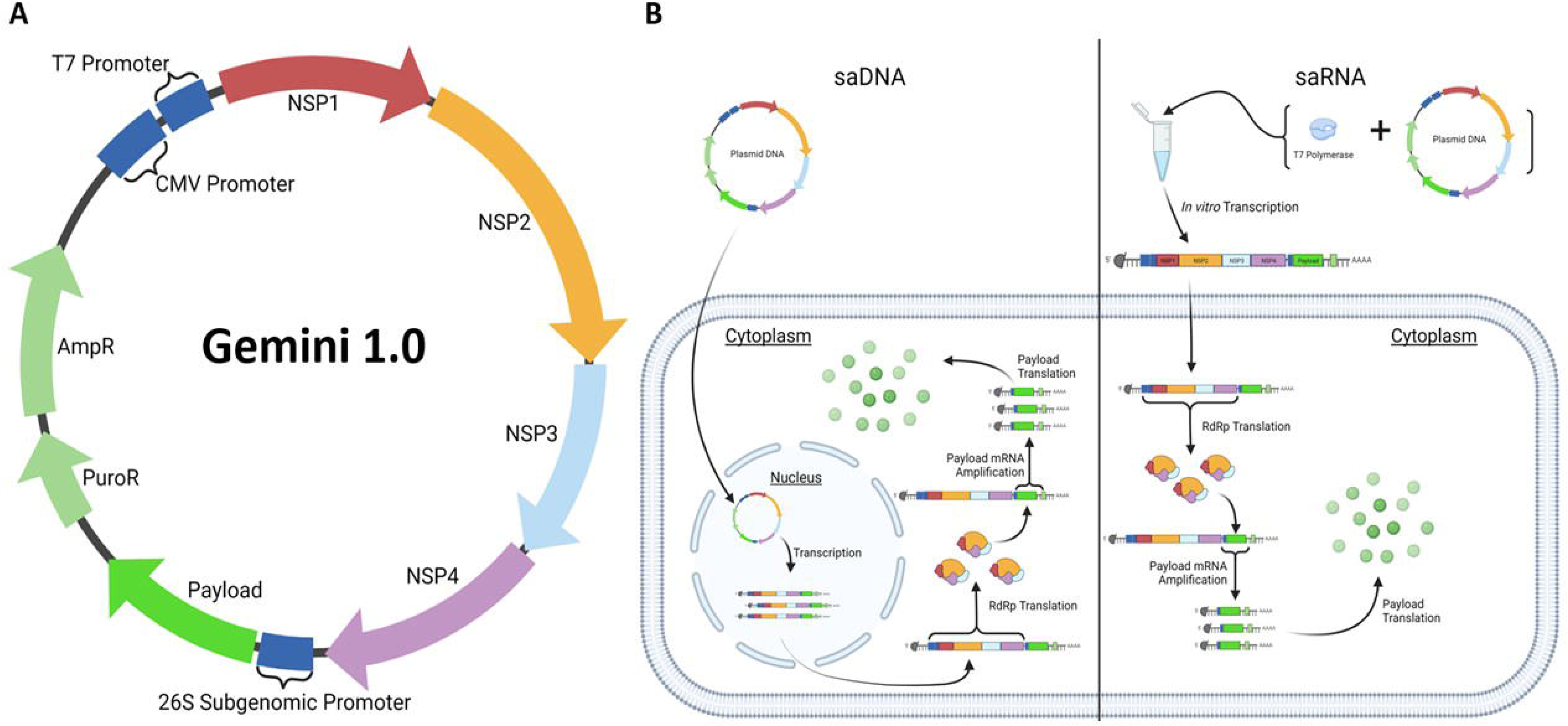
**(A) Genomic map of the Gemini 1.0 vector**. The vector contains a binary promoter (CMV and T7), replicon protein genes (NSP1-4), a 26S sub-genomic promoter from the Venezuelan equine encephalitis (VEE) virus, and resistance genes for puromycin (PuroR) and ampicillin (AmpR) for mammalian and bacterial cell culture, respectively. **(B) Delivery mechanisms of Gemini as saDNA (Gemini-D) or saRNA (Gemini-R). In-vitro replication**. Gemini-D (left) is delivered directly into the cell; it enters the nucleus for transcription then re-enters the cytoplasm for translation using the endogenous machinery to express its RNA dependent RNA polymerase (RdRp). Gemini-R (right) is delivered into the cytoplasm for direct translation of its RdRp by the endogenous machinery. The RdRp then transcribes the payload mRNA, allowing for production of the payload protein. saRNA is transcribed in vitro (top right) via the T7 promoter with the assistance of T7 polymerase. Also, 5’ capping is performed before mRNA delivery into the cytoplasm for direct translation of RdRp and subsequent expression of payload mRNA.

**Fig. 2.**
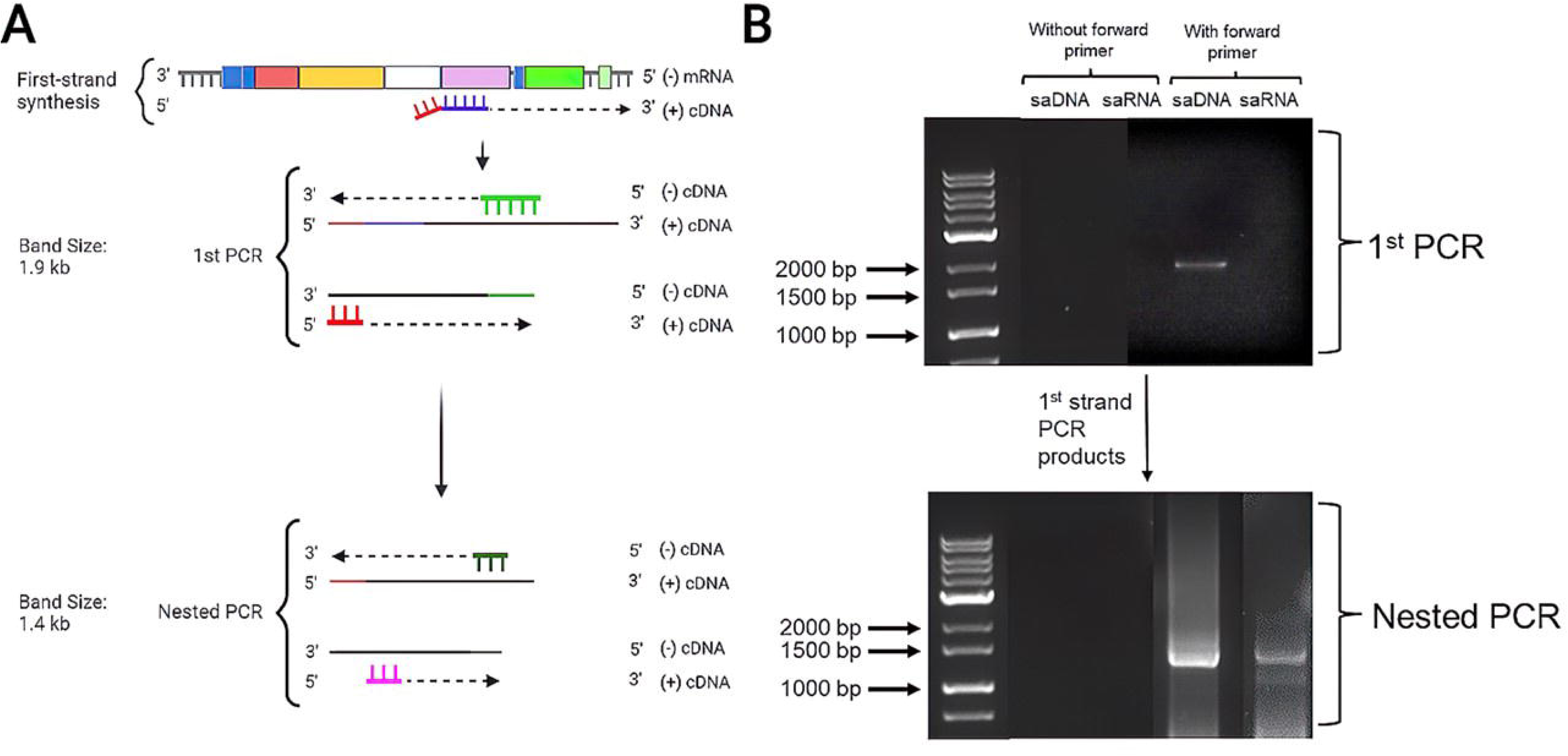
Negative-strand RNA is present post-transfection with eGFP Gemini-R and Gemini-D. **(A)** Detection strategy: First-strand synthesis is conducted using a NSP4 region specific primer (purple) with a random nucleotide tag (red) to produce positive strand (+) cDNA. The negative (-) RNA strand is then synthesized as cDNA using primers specific for the eGFP region (green) and the random nucleotide tag, producing a band of about 1.9 kb. A subsequent nested PCR (primers: dark green and fuchsia) then produces a band of 1.4 kb. **(B)** Results of applying the strategy described in panel (A) to yield total RNA from HEK293 cells transfected with Gemini-R and Gemini-D expressing eGFP. Results of the first and nested PCRs are shown when the forward primer was used (columns 4 and 5) and not used (columns 2 and 3, control). The reference ladder in column 1 indicates the position and sizes of the molecular weight markers in base pairs (bp).

PCR products of the expected size were detected as the nested PCR product of total RNA extracted from transfected cells with either Gemini-R or Gemini-D (Figure 2B). Their sample source can be attributed to the presence of the PCR product from the first round of PCR. To verify the fidelity of the platform, it was determined that the production of the requisite negative (-) RNA strand from the transfected cells was not attributable to primer-independent effects, it was observed that when a gene-specific forward primer was omitted during cDNA synthesis, no PCR products were present in either the first PCR or in the subsequent nested PCR (Figure 2B).

### Both Gemini-R and Gemini-D drive protein expression in transfected cells

Validation that Gemini is capable of driving expression of a clinically relevant payload from either format was conducted by transfecting HEK293 cells with either Gemini-R or Gemini-D, both expressing spike protein from the B.1.617.2 (Delta) variant of the SARS-CoV-2 virus.

Western Blot analysis of protein expression at day 6 after transfection confirmed the presence of protein bands of the expected size (Figure 3A). Western Blot results were subsequently reinforced by a flow-cytometric analysis of expression in transfected cells carried out over a time-course of 6 days. In the absence of any selection, by day 2 post-transfection, the Gemini-D driven form of Spike protein expressed in HEK293 cells revealed a weak positivity for spike protein expression (3.85% of the total cell population), but by day 6 the positive fraction had significantly increased (24.0% of the total population), suggesting self-amplification behaviour of Gemini-D (Figure 3B-3D). Similarly, by day 2 post-transfection, the Gemini-R driven form of Spike protein expressed in HEK293 cells revealed a weak positivity for spike protein expression (3.70% of the total cell population), but by day 6 the positive fraction had significantly increased (31.6% of the total cell population), suggesting self-amplification behaviour of Gemini-R (Figure 3B-3D). In contrast, cells transfected with a conventional LNP-encapsulated pseudouridine substitute mRNA encoding the B.1.617.2 (Delta) spike protein variant of the SARS-CoV-2 virus were 36.0% positive for the spike protein initially on day 2 post-transfection, but by day 6, there was a 6-fold reduction and the majority of the transfected cells had become negative (6.08%; Figure 3B); demonstrating a rapid waning of expression *in vitro*.

**Fig. 3.**
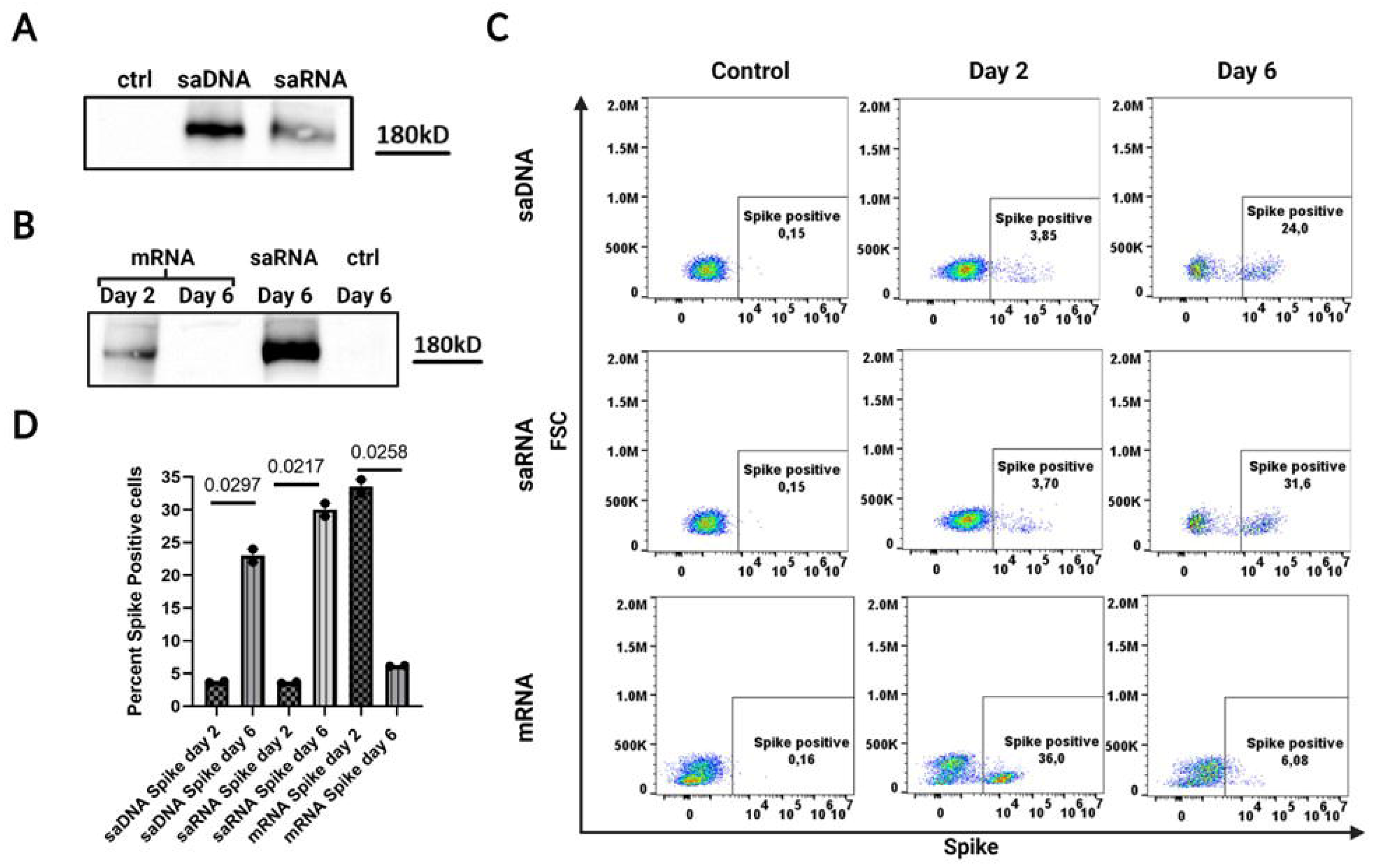
Temporal dynamics of protein expression in HEK293 cells transfected with Gemini-R and Gemini-D expressing SARS-CoV-2 spike protein. LNP-encapsulated B.1.617.2 (Delta) spike protein variant of the SARS-CoV-2 virus expressed in saDNA (Gemini-D) or saRNA (Gemini-R) in HEK293 cells were compared with a conventional LNP-encapsulated plasmid DNA vector expressing the B.1.617.2 (Delta) spike protein variant of the SARS-CoV-2 virus on day 2 and 6 using unpaired t-test. **(A)** Western Blot at day 6 after transfection **(B)** mRNA expression over time **(C)** Results of flow cytometry. Spike-expressing cells were subject to flow cytometric analysis on Day 2 and 6 post-transfection, using a BD Cytoflex flow cytometer **(D)** Quantitated fraction of positive cells (in percent’s). p-values are indicated.

### eGFP expression induced by Gemini-D and Gemini-R in transfected cells and in injected mice

To compare *in vitro* payload expression capabilities of the saDNA and saRNA platform, HEK293 cells were transfected with either Gemini-R or Gemini-D expressing eGFP. Western blot analysis confirmed protein expression by yielding a protein band of the expected size at day 6 after transfection and sorting (Figure 4A). Two days post-transfection, cells were cell-sorted to obtain 100% eGFP positive cells. A flow-cytometric analysis of such cells every week up to 6 weeks were performed (Figure 4A, Figure 4C). We found cells transfected with Gemini-D to be strongly positive on day 14 (98%). By day 28, this positive fraction had only marginally decreased to 93%, suggesting long-term expression from the Gemini-D self-amplificating format. In slight contrast, cells transfected with Gemini-R were 85% positive on day 14, and, by day 28, the corresponding fraction had significantly dropped to 64%. This suggests that expression from the Gemini-D is more stable than Gemini-R in transfected cells; the difference in the fraction of positive cells is statistically significant at both 14- and 28-days post-transfection (Figure 4D) (*p*-value of unpaired *t*-test are 0.0007 and 0.0001, respectively). In contrast cells transfected and sorted for eGFP expressing a conventional LNP-encapsulated pseudouridine substitute mRNA encoding eGFP are initially most positive for eGFP (92.6%) but by day 6 (Figure 4C) there is a rapid reduction in protein expression as detected by Western blotting (Figure 4B), and nearly a 3-fold reduction in detectable eGFP expressing cells (28.2%) detected by flow cytometry (Figure 4C). The *p*-value of unpaired *t*-test between day 2 and 6 is 0.0001 (Figure 4D).

**Fig. 4.**
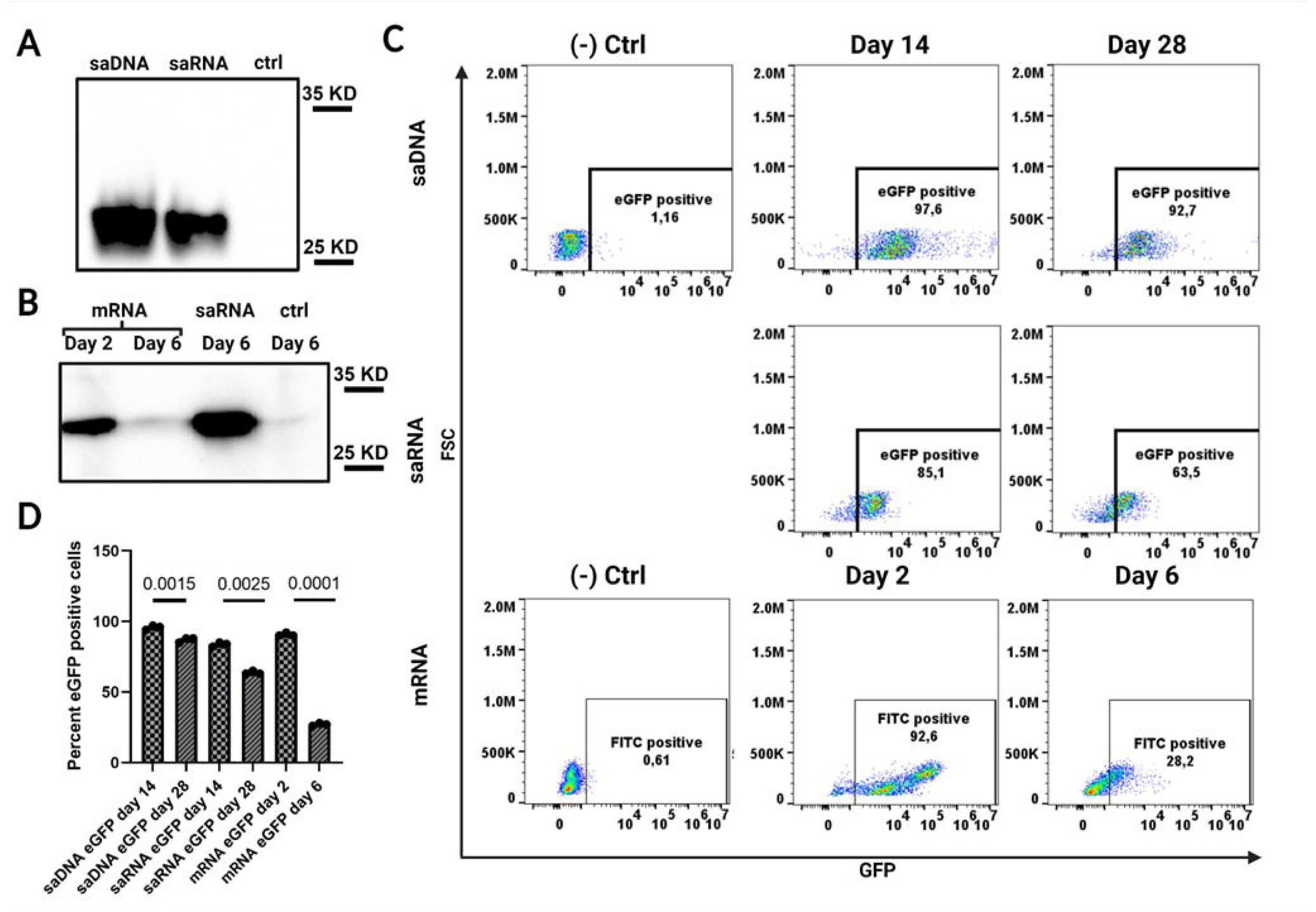
Temporal dynamics of protein expression in HEK293 cells transfected with Gemini-R and Gemini-D expressing eGFP protein. Time course of LNP-encapsulated eGFP positive saDNA (Gemini-D) and saRNA (Gemini-R) cells were compared on weeks 2 and 4 after selection by cell sorting for eGFP positive. **(A)** Western Blot at day 6 after sorting **(B)** mRNA expression over time **(C)** HEK-293 cells were transfected with Gemini-D and Gemini-R expressing eGFP and the samples were sorted for GFP expression 2 days post-transfection. Sorted GFP-expressing cells were then subject to flow cytometric analysis on Day 2, then weekly up to 6 weeks for EGFP expression using a BD Cytoflex flow cytometer **(D)** Quantitated fraction of positive cells (in percent). eGFP positive saDNA and saRNA cells were compared on weeks 2 and 4 using unpaired t-test. p-values are indicated.

To evaluate such expression dynamics *in vivo*, mice were intramuscularly injected with the same Gemini-R or Gemini-D encoding eGFP. Injected leg muscles were collected at three different time points (14-, 28- or 42-days post-injection, n=3 per day of sacrifice), frozen and subsequently sectioned (as described in the Materials and Methods), then assessed for native eGFP expression both qualitatively (Figure 5A) and quantitatively (Figure 5B, 5C). Compared to the negative control, all time points except the day 14 for Gemini-D and day 42 for Gemini-R are less than p<0.05. For Gemini-R, high levels of eGFP were observed 14 days post injection, with levels appearing reduced at subsequent time points. On the other hand, expression levels from Gemini-D continued increasing and appeared highest at the latest time point, 42 days after injection.

**Fig. 5.**
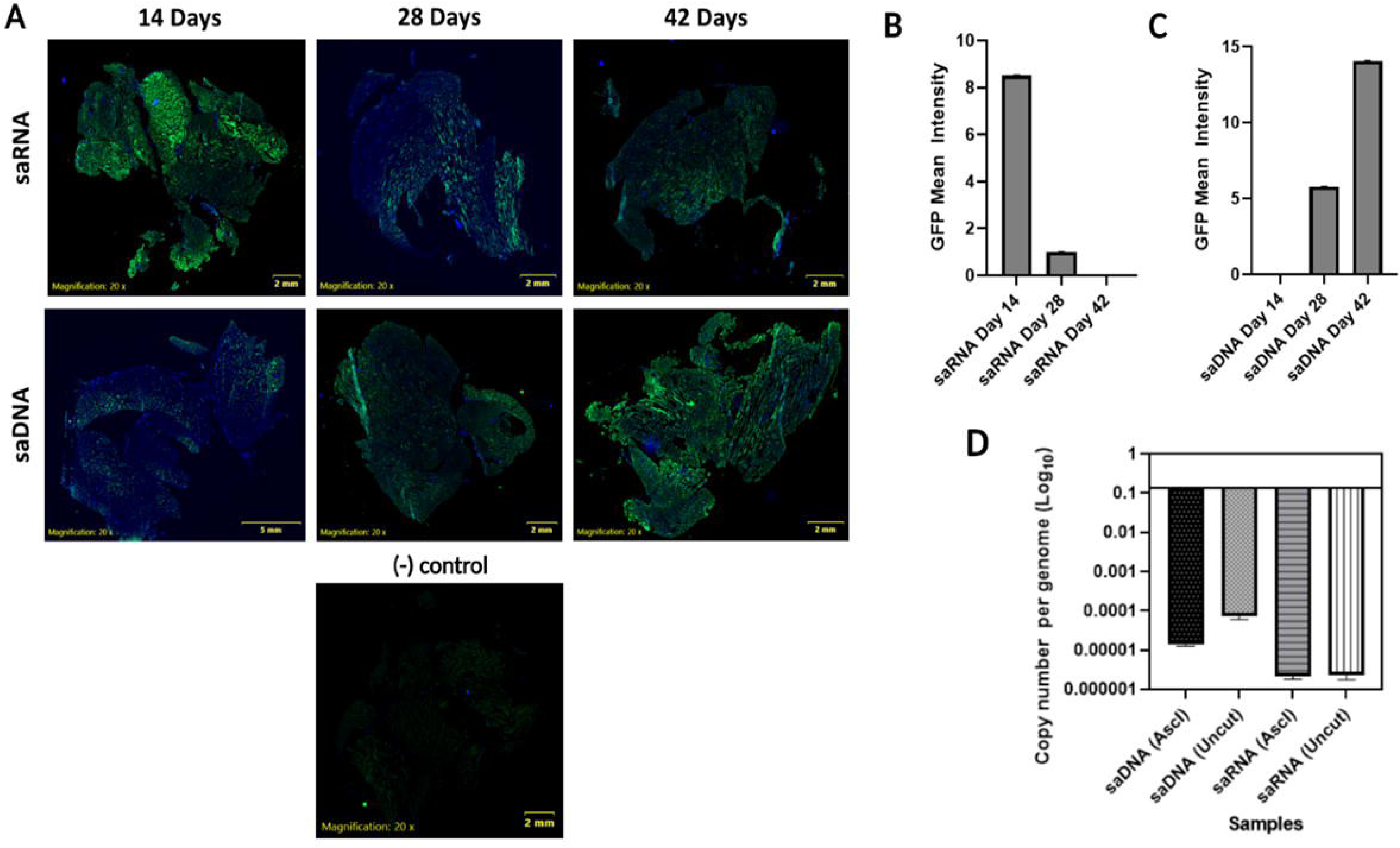
Gemini-R and Gemini-D driven eGFP expression and genome integration *in vivo*. Both Gemini-R and Gemini-D achieve long lasting expression of a protein payload in vivo and exhibit negligible genome integration. **(A)** eGFP fluorescence induced by Gemini-R and Gemini-D in mouse muscle tissue after 14, 28 and 42 days. **(B)** Quantitative fluorescence for Gemini-R estimated from panel (A). All time points except the day 14 for Gemini-D and day 42 for Gemini-R are less than p<0.05, compared to the negative control. **(C)** Quantitative fluorescence for Gemini-D estimated from panel (A). **(D)** Estimated frequency of integration of Gemini-R and Gemini-D in the host genome (see Materials and Methods).

Overall, the expression of eGFP from Gemini-R and Gemini-D has been demonstrated to persist for over 28 days, with Gemini-D being still highly active and potentially increasing in expression *in vivo* after 42 days while gene expression using conventional mRNAs diminish after a few days. The results with Gemini are consistent with the duration of gene expression achieved by viral vector vaccine platforms such as the ones based on recombinant adenoviruses or vaccinia virus (13, 14). Thus, though Gemini-D is itself is a non-viral based plasmid format, its self-amplifying capacity allows it to last some 40 days *in vivo*. The data suggest that Gemini-R may be useful for delivery and expression of therapeutic payloads for 4-weeks while Gemini-D may be useful for applications that require longer periods of expression.

### Neither Gemini-R nor Gemini-D induce significant genomic integration

We utilized gel-based methods to separate genomic DNA from extrachromosomal DNA (15) and the properties of restriction enzyme AscIS to detect genomic integration in muscle (16).

While it cleaves DNA derived from prokaryotic cells, AscIS selectively ignores restriction sites that are CpG methylated in eukaryotic cells:

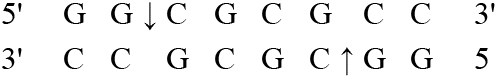

We used it and established agarose gel procedure to distinguish between integrated and extrachromosomal nucleic acids to estimate the *in vivo* frequency of Gemini-R and Gemini-D integration in the genome of mouse muscle cells (see Materials and Methods).

While the Gemini-R format showed no detectable integration in the muscle injection site, the number of Gemini-D integrated copies was determined to be less than 3×10^−6^ per cell genome (see Figure 5D). Such integration rate in muscle tissue is significantly lower than that of plasmid DNA, 5×10^−5^(17), or adenoviruses, 6.7×10^−5^ (18). As recombinant adenoviruses are widely used clinically and considered one of the safest vaccine vectors available (18), this result demonstrates an excellent safety profile for Gemini-D and allays fears of induced genetic abnormalities, such as those resulting from lentiviruses and other commonly used retroviral expression systems (More detailed comparisons are shown in Supplementary Table 1S).

Thus, overall, Gemini-R was demonstrated not to integrate into the host genome while Gemini-D was determined to be less than 3×10^−6^ per cell genome, lower than other clinically approved vector platforms and many orders of magnitude lower than the spontaneous somatic mutation frequency (Supplementary Table 1) in mice or humans (19, 20) establishing this as one of the safest platforms yet created.

### Significant Antibody response are induced by both SARS-CoV-2 spike Gemini-R and Gemini-D vaccine formats

The antibody response derived against a relevant vaccine payload was evaluated in mice. Gemini-R and Gemini-D vectors expressing the B.1.617.2 (Delta) spike variant of the SARS-CoV-2 virus were LNP-encapsulated (see Materials and Methods) and 5ug (the optimal dose) of either vaccine formulation was injected into K18-hACE2 transgenic mice expressing the human ACE-2 SARS-CoV-2 receptor according to the vaccination protocol outlined in materials and methods. All the treatment groups were statistically different than the unvaccinated control group (ANOVA, *p* = 0.0082). Consistent with the flow cytometry findings described above, the IgG response to the Gemini-D format of the vaccine in serum extracted from inoculated mice on day 28 (Figure 6B) was found to be significantly greater (*p* = 0.0046) when compared to the unvaccinated control group, while the IgG response to the Gemini-R format of the vaccine was found to be lower than that achieved with the Gemini-D format (*p* = 0.1019) compared to the unvaccinated control group. In contrast, the IgM response (Figure 6A) demonstrated opposite behavior, with significantly higher IgM levels generated by the Gemini-R format of the vaccine (*p* = 0.0420; here the R-Gemini group was compared to the unvaccinated group with unvaccinated background being subtracted from both the groups), and a lower IgM response generated by the Gemini-D format of the vaccine (*p* = 0.1525, here the D-Gemini group was compared to the unvaccinated group with unvaccinated background being subtracted from both the groups). Finally, neither vaccine format induced a significant IgA antibody response nor any observable toxicity resulting in weight loss, a parameter of safety and toxicity in mice (p > 0.05, data not shown).

**Fig. 6.**
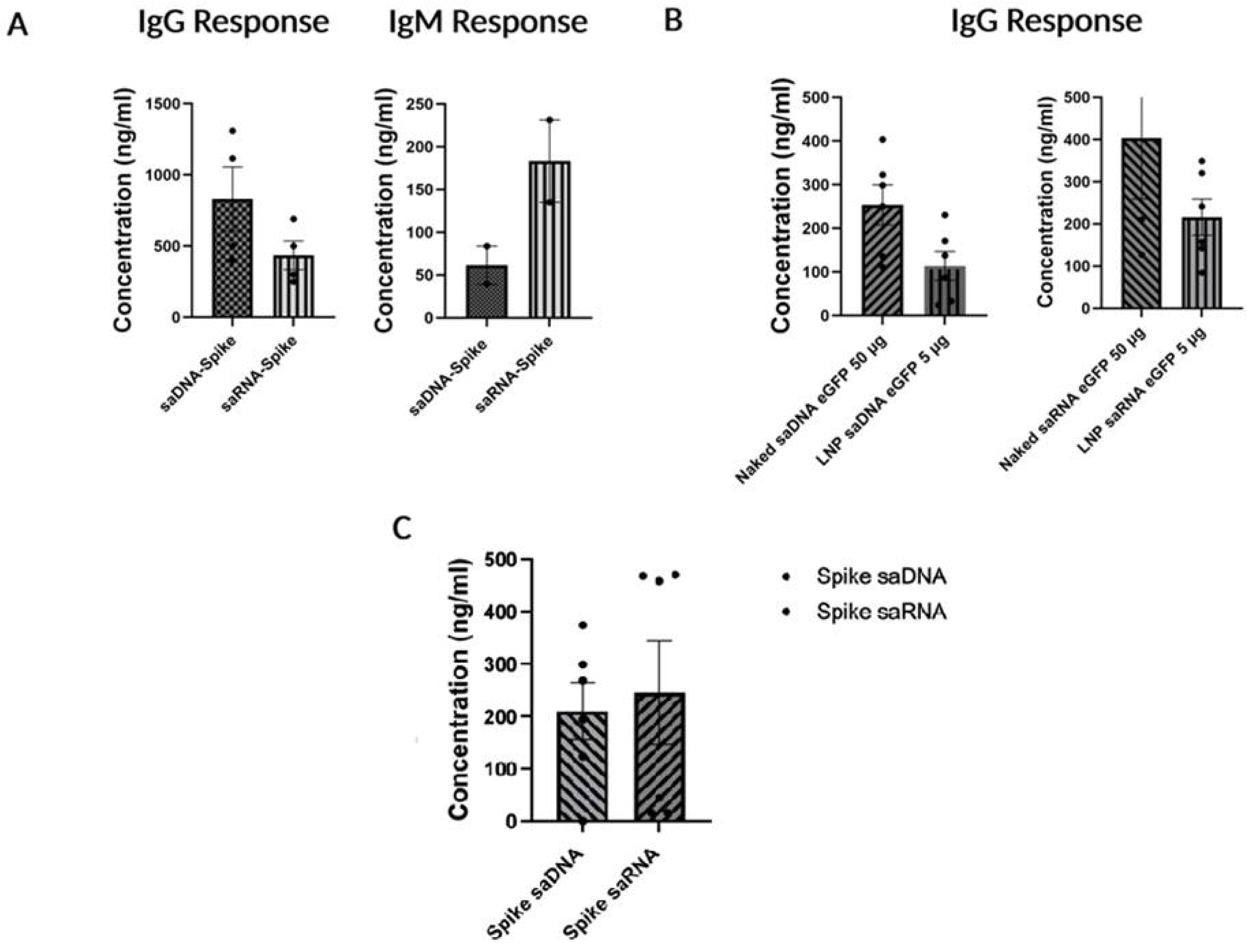
Immune response in mice inoculated with Gemini-D and Gemini-R expressing SARS-CoV-2 spike protein. A single dose of either the LNP-encapsulated Gemini-R and Gemini-D vaccine formats elicits significant antibody concentrations **(A)** Antibody concentration measured in mouse serum 28 days post-injection: Left: IgG response; Right: IgM response. All the treatment groups were statistically different than the unvaccinated control group (ANOVA, *p* = 0.0082). **(B)** Comparison between immune response to naked and encapsulated eGFP: Left: Gemini-D; Right: Gemini-R. All the treatment groups were statistically different than the unvaccinated control group (ANOVA, *p* = 0.0134). See Materials and Methods for the meaning of bars and points. **(C)** Immune response in mice inoculated with naked formulation of Gemini-D and Gemini-R expressing Omicron SARS-CoV-2 spike protein. A single dose of either the naked Gemini-R and Gemini-D encoding the B.1.1.529 (Omicron) Spike variant of SARS CoV-2 Spike vaccine format elicits significant cross-reactive antibody concentrations against B.1.617.2 (Delta) spike variant of the SARS-CoV-2 virus. Mice were inoculated on day 0 with Naked (saDNA) Gemini-D (50 µg) and (saRNA) Gemini-R (50 µg) intramuscularly. The sera were collected from all the groups measured at 42 days post-vaccination and SARS-CoV-2 spike-specific IgG levels were assessed using indirect B.1.617.2 (Delta) spike variant of the SARS-CoV-2 virus ELISA. All values are less then p <0.05 for vaccinated groups compared to normal saline subtracted from the treated groups.

Comparable absolute quantitative estimates in the clinical literature are limited, as most estimates are based on international units. However, the few available sources of absolute quantitation of antibody responses such as (21), where they tested several thousand patients, support the findings that either LNP-encapsulated Gemini-R and Gemini-D format elicit robust antibody responses; in quantitative terms (ng/ml). Overall, a single dose of 5ug of either the LNP-encapsulated Gemini-R and Gemini-D formats (Figure 6B) elicit significantly greater IgG concentrations (400-800 ng/ml) compared to those observed in patients mounting immune responses to SARS CoV-2, which on average reach ∼200 ng/ml. Finally, following intramuscular injections, Gemini-R may have advantages in vaccines where IgM has the greatest clinical value while Gemini-D may find its greatest use in applications where immunity to specific pathogens require IgG responses. It is conceivable that these differences may be due to differential Toll-like receptor (TLR) signally pathway for RNA and DNA (22). Thus, utilizing different Gemini formats may provide the opportunity to “tune” immune responses towards an IgG or IgM dominant immune response.

### LNP encapsulation is not necessary for the immunogenicity of either Gemini-D- or Gemini-R vaccine formats

Current paradigms for RNA delivery require encapsulation of either mRNA or saRNA through expensive and technically demanding formulations. In contrast to this, protecting saRNA from RNAse digestion, either on the interior formulated LNP or on the exterior of pre-made cationic LNP particles (23) protects saRNA from RNAse digestion and, after vaccination, induces a statistically equivalent amount of antibodies against the HIV-1 Env gp140 protein used as a model antigen (23).

In the context of this study, we thought it prudent to addressed whether LNP encapsulation is necessary for successful delivery and performance of the vaccines based of either Gemini-D or Gemini-R using eGFP as a “model antigen”. All the treatment groups were statistically different than the unvaccinated control group (ANOVA, *p* = 0.0134). Gemini vaccines expressing eGFP were tested (Figure 6C) and showed that higher doses (50ug) of naked Gemini-D performs better (*p* = 0.0129) when compared with 5ug of LNP-encapsulated Gemini-D (*p* = 0.1983). We achieved similar results were consistent for 50ug of naked (*p* = 0.0420) and 5ug of encapsulated Gemini-R (*p* = 0.3269) expressing eGFP as a model antigen when compared with control groups. Vaccination of mice with 50ug of either naked Gemini-R or Gemini-D encoding the B.1.1.529 (Omicron) Spike variant of SARS CoV-2 Spike also elicits a robust immune response that is produces antibodies cross-reactive with the B.1.617.2 (Delta) spike variant of the SARS-CoV-2 virus (Figure 6C). These data imply that dosing alone can overcome the need for LNP encapsulation.

The findings in this study seemingly contradict the accepted wisdom on previously described saRNA platforms (23) by offering the encouraging possibility that LNPs may be omitted altogether in future saRNA-based vaccines and simple dosing might be used to dispense with the technically demanding LNP encapsulation of vaccine payloads delivered through either Gemini-R and Gemini-D. Furthermore, the implications are particularly important for Gemini-D saDNA-based vaccine format, which can be rapidly prepared as a simple plasmid without the need for LNPs of any kind.

## Conclusions and outlook

Our study demonstrates that the Gemini platform is useful for the creation of recombinant vaccines and potentially other payloads that may be of use therapeutically and provides several advantages when compared with other platforms such as the conventional recombinant mRNA and DNA technologies.

There are clear advantages of the dual format of Gemini, as it safely combines the flexibility and power of RNA platforms with the much greater stability and ease of manufacturing of DNA constructs. In particular, the self-amplifying DNA plasmid format used in the Gemini-D format can be very rapidly created, scaled-up in very large conventional manufacturing batches, resulting in better standardization, without being impeded by the production bottlenecks incurred by the highly technical manufacturing procedures required for conventional RNA vaccines; this favorable property would assure a swifter response than currently imagined to a future pandemic. Gemini-D is similarly convenient in terms of distribution and thermal stability storage as, unlike mRNA vectors, it is stable, and it does not need ultra-low storage temperatures during transportation. Furthermore, the dual expression platform offers the ability to choose and directly compare either a saRNA or saDNA platform while retaining the same payloads. Thus, Gemini may overcome issues associated with vaccine stability, attributed to the requirements of prolonged ultra-low temperature storage, avoiding logistical and practical concerns associated with the world-wide distribution of vaccines (24).

In this study, the B.1.617.2 (Delta) spike protein variant of the SARS-CoV-2 virus and the B.1.1.529 (Omicron) Spike variant of SARS CoV-2 Spike and the eGFP gene were chosen as “model” antigenic payloads to establish the proof-of-concept of both the Gemini-D and Gemini-R due to the current interest in SARS-CoV-2 vaccines and the established utility of eGFP as faithful reporter protein antigen. Both Gemini-R and Gemini-D are able to express payload proteins for over 28 days *in vivo* in mouse muscle, which is similar to the duration achieved by viral expression vectors such as those based on adenoviruses or vaccinia virus.

Furthermore, it should be noted that transfection of HEK293 cells with a conventional LNP-pseudouridine substituted mRNA analogous to the Moderna vaccine intitiates expression by 2 days but the expression of spike protein from the B.1.617.2 (Delta) variant of the SARS-CoV-2 virus or eGFP diminishes 3-6 fold by only 6-7 days of transfected cells. Early studies on the initiation of humour immunity by B-lymphocytes established that lower antibody responses were noted in animals receiving exposure to antigen for less than 4 days (21). Subsequent studies on T-lymphocyte responses to viruses demonstrated that a similar minimum duration of temporal exposure to the antigen of 4 days was necessary to generate maximal cell-mediated immunity and immunological T-lymphocyte memory (22). Our *in vitro* work demonstrates that self-limiting conventional LNP-mRNA used in widely distributed SARS-CoV-2 vaccines could result in rapidly waning antigen expression *in vivo* with a timescale of a few days. Therefore, it is an open question whether the transient antigen expression provided by such delivery systems might be related to the decline in immunity observed after only 3 months following vaccination against SARS-CoV-2 with LNP-mRNA (23).

In addition, a key and elegant innovation in the recent mRNA vaccine revolution was the inclusion of pseudouridines during transcription to create a highly expressed, non-immunogenic, non-inflammatory platform (25). Paradoxically, the LNPs used to deliver the mRNA vaccines have been found to be highly pro-inflammatory and thus, may contribute to some of the observed side effects of these LNP-encapsulated vaccines (26). Both Gemini platforms are efficacious without the need for LNP-encapsulated and therefore reduce the potential for adverse reactions due to the noted highly inflammatory nature of LNPs (26).

Taken together, the Gemini platform possesses attractive properties with respect to storage and safety profiles that likely exceed other recombinant vaccine platforms, while eliminating the need for LNP encapsulation. Finally, it is impactful that non-encapsulated Gemini-D, the self-amplifying DNA plasmid format, is simple enough to be made in laboratories with limited technical resources. In the longer term, the Gemini platform may lead to a true democratization in the creation, manufacturing, and distribution of vaccines and therapeutics.

## Materials and Methods

### Vector synthesis

Figure 1A illustrates the map of the vector system discussed in this paper, hereinafter referred to as Gemini 1.0. It is based on the T7-VEE-GFP plasmid a very gift from Professor Steven Dowdy, at Department of Cellular & Molecular Medicine, University of California, San Diego School of Medicine, 9500 Gilman Drive, La Jolla, CA 92093-0686, USA (27). It consists of the NSP1-4 genes from Venezuelan equine encephalitis (VEE) virus, an origin of replication site, a bacterial promoter (26S subgenomic promoter), an Ampicillin resistance (AmpR) gene acting as a selection marker for bacterial culture, a T7 promoter to recruit T7 RNA polymerase for saRNA synthesis, and a human CMV enhancer/promoter, for use as a DNA or RNA vector in humans. The CMV promoter was subsequently cloned into the T7-VEE-GFP plasmid by Synbio Technologies. For the B.1.1.529 (Omicron) Spike variant of SARS CoV-2 and B.1.617.2 (Delta) spike variant of the SARS-CoV-2 virus and eGFP sequences see Appendix 1.

### saDNA and saRNA preparation, conventional Plasmid DNA and mRNAs

The Gemini 1.0 vector was transformed into DH5α Competent *E. Coli* (NEB, C2987) and plated onto Luria-Bertani (LB) agar containing Ampicillin for selection; this was followed by overnight culturing in LB broth at 37 °C. Plasmid DNA was extracted according to the EZ10 Plasmid DNA Minipreps Kit protocol (BioBasic, BS6149). To prepare the saRNA, the Gemini 1.0 plasmid underwent in vitro transcription using T7 RNA polymerase (NEB, M0251L), followed by in vitro 5’ capping and 3’ polyadenylation. Figure 1 describes the self-amplifying platform pathways and *in vitro* replication process for both the DNA and RNA forms.

### mRNA preparation

Pseudouridine substitute LNP-encapsulated mRNAs encoding eGFP (Cat # PM-LNP-21) or LNP-B.1.617.2 (Delta) spike protein variant of the SARS-CoV-2 virus (Cat # PM-LNP-12) mRNA purchased from Promab Biotechnologies, 2600 Hilltop Dr Building B, Suite C320, Richmond, CA 94806, United States. These LNP’s were formulated by Prolab with, SM-102, DSPC, cholesterol, and DMG-PEG2000 at optimal molar concentration for a high rate of encapsulation and efficient mRNA delivery.

### Lipid Nanoparticles (LNPs) formulation of Gemini-D and Gemini-R formats

LNP-encapsulated forms of D- and Gemini-R were prepared by mixing 5 µg of saDNA or saRNA with 18 µL of Genesome lipid solution (DOTAP:Chol:DOPE in a 1:0.75:0.5 ratio; Encapsula Nano Science, GEN-7036) in a 1:2 volume ratio at room temperature as described by the manufacturer. LNP-protected nucleic acids were kept on ice until ready for injection.

### HEK293 cell culture and transfection with Gemini

HEK293 cells (ATCC;CRL-1573) were cultured in Dulbecco’s Modified Eagle Medium (DMEM; Gibco, 11965-092) containing 10% Fetal Bovine Serum (FBS; Gibco, A3160702) and penicillin/streptomycin. Cells were seeded at a density of 5*105 cells per well in a 6-well plate one day prior to transfection. Transfection was performed with ∼ 2.5 µg of either D- or Gemini-R according to the protocol for LipofectamineTM 3000 (ThermoFisher Scientific, L3000001).

### Negative (-) strand mRNA detection

HEK293 cells were harvested 72 hrs post-transfection. Total RNA was extracted using the PureLinkTM RNA Mini Kit (Ambion, 12183025) and its integrity was checked on a 0.8% agarose gel. Thereafter, total RNA was treated with amplification grade DNase I (Invitrogen, 18068015) to remove any residual DNA, followed by first strand cDNA synthesis using either a NSP4 gene-specific forward primer with a random nucleotide tag sequence (5’-cggtcatggtggcgaataaGCGGCCTTTAATGTGGAATG-3) or without any primer according to the SuperScriptTM III Reverse Transcriptase protocol (Invitrogen, 18080044). cDNA synthesis was then completed followed by a PCR using an eGFP gene-specific reverse primer (5’-CACCTTGATGCCGTTCTTCT-3’) and the random nucleotide tag-specific forward primer (5’-cggtcatggtggcgaataa-3’) to produce a 1.9 kb band which would be an indication of negative RNA strand. A nested PCR using the forward primer, 5’-CCGAGAGCTGGTTAGGAGATTA-3’, and reverse primer, 5’-GCTTGTCGGCCATGATATAGA-3’ on the first PCR product was then performed to amplify cDNA with a band size of 1.4 kb to verify the correct target sequence (see Figure 2A). Parameters used for both PCRs: 94 °C for 30 seconds, 56 °C for 30 seconds, 72 °C for 30 seconds, for 28 cycles.

### Flow cytometry sample preparation

HEK293 Cells were transfected with either: (1) Gemini-D expressing SARS-CoV-2 spike protein, or (2) a non-self-amplifying DNA plasmid control expressing SARS-CoV-2 spike (see Figure 2A-ii) (see Appendix for the sequences) or LNP-B.1.617.2 (Delta) spike protein variant of the SARS-CoV-2 virus mRNA or the conventional LNP-encapsulated Pseudouridine substitute mRNAs encoding the B.1.617.2 (Delta) spike protein variant of the SARS-CoV-2 virus (Appendix Fig S1-S3). Cells were harvested at 2- and 6-days post-transfection as were the FACS samples and resuspended in FACS buffer at a concentration of 1.0×10^6^ cells/mL. Subsequently, cells were incubated with FACS buffer for 30 minutes at 4 °C cells for Fc blocking, centrifuged (1200 RPM for 4 minutes at 4 °C), and then stained with an anti-receptor-binding domain (RBD) of SARS-CoV-2 spike antibody conjugated to Alexa Fluor 647 (1:100; invitrogen, 51-6490-82) for 30 minutes at 4 °C away from light. Cells were then washed in FACS buffer (1200 RPM for 4 minutes at 4 °C) and resuspended in FACS buffer (500 uL/tube). They were prepared similarly to the FACS samples in the following section, the data was acquired using BD Cytoflex flow cytometer.

### Fluorescence-activated cell sorting (FACS) sample preparation

HEK293 Cells were transfected with either a: (1) Gemini-D expressing eGFP, (2) Gemini-R expressing eGFP, or (3) LNP-encapsulated Pseudouridine substitute mRNAs encoding eGFP (Cat # PM-LNP-21) mRNA. (See Appendix 1 for the sequences). Two days post-transfection, cells were sorted for eGFP expression. Cells were then harvested at 14- and 28-days post-transfection using Cellstripper® (Corning, 25-056-CI), counted using a TC20 Automated Cell Counter (Bio-Rad, 1450102), and resuspended in FACS buffer (1X phosphate buffered saline (PBS) with 2% FBS and 2% normal rabbit serum) at a concentration of 1.0×10^6^ cells/mL. Subsequently, cells were incubated with FACS buffer for 30 minutes at 4 °C cells for Fc blocking, centrifuged (1200 RPM for 4 minutes at 4 °C), and resuspended in FACS buffer (500 uL/tube). Data was acquired using a BD Cytoflex flow cytometer.

### Western blot

HEK293 cells were transfected with either: (1) D- or Gemini-R expressing SARS-CoV-2 spike (2) D- or LNP-encapsulated Pseudouridine substitute mRNAs encoding eGFP (Cat # PM-LNP-21) or LNP-B.1.617.2 (Delta) spike protein variant of the SARS-CoV-2 virus (Cat # PM-LNP-12) mRNA (See Appendix 1). Cells were harvested 72 hours post-transfection, lysed in 2X sample buffer supplemented with 5% β-Mercaptoethanol (BME; Bio-Rad, 1610710), and heated at 90 °C for 10 minutes. Subsequently, samples were treated with Benzonase nuclease (Sigma, E1014) for 3 hours to remove nucleic acids. A total of 30 µg of protein per well was loaded onto a 4-15% precast SDS-PAGE gel (Bio-Rad, 4561083). SDS-PAGE running conditions are as follows: 75V for 20 minutes, then 120V for 2 hours. Protein was transferred to a nitrocellulose membrane using 75V for 3 hours. Subsequently, the membrane was washed in 0.1% Tween-20 in PBS (PBST) followed by blocking with 5% skim milk in 1% PBST for 2 hours. Primary antibodies for ALFA Tag (Nano-tech, N1581) and eGFP (UBC AbLab, 21-0024-01) were diluted 1:5000 and incubated with the membrane at 4 °C overnight. The membrane was then treated with three 10 minutes washes with 0.1% PBST. Anti-rabbit (ThermoFisher Scientific, 31460) and anti-mouse (Abcam, AB205719) secondary antibodies conjugated to HRP were diluted 1:5000 and incubated with the membrane at room temperature for 1 hour followed by three 10-minute washes of 0.1% PBST. Subsequent signal detection was conducted on a ChemiDoc Imaging System (Bio-Rad).

### Qualitative determination of eGFP expression in injected mouse leg muscles

Two groups of 6–12-week-old K18-hACE2 mice were injected with 5 ug of either LNP-Gemini-R or LNP-Gemini-D expressing eGFP (see Appendix Figure S1 and S2 for the sequence), by intramuscular injection into the caudal thigh muscle. For each vaccine group, mice were sacrificed, at 14-, 28- or 42-days post-injection (n=3 per day of sacrifice). Thigh muscles were excised and immediately frozen on dry ice in Neg-50™ (Richard-Allen Scientific, Thermo Scientific). Samples were stored at 80 °C until they were sectioned, using a cryostat microtome, and counter stained with DAPI at the Centre for Phenogenomics, University of Toronto. Images were captured at the same facility (Figure 4) and were subsequently assessed for qualitative expression of native eGFP. Four sections per sample were assessed; the one with highest eGFP intensity was chosen per sample. A visual ‘average’ was ascertained from these images for each time point and a suitable representative image was selected.

The highest eGFP intensity images were used to quantify eGFP mean intensity (Figures 4B-Figure 4D), utilizing ImageJ software to calculate mean intensity and total area for each sample. Weighted mean intensities were calculated for each vaccine group per time point, as the sum of the individual weighted mean eGFP intensities per sample. The individual weighted means were calculated using the following equation:

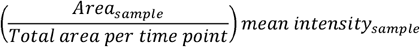

### Immunization of mice

K18-hACE2 transgenic mice were purchased from the Jackson Laboratory and maintained in the Centre for Disease Modeling at the University of British Columbia. These experiments were approved by the Animal Care Committee (UBC). Animals were maintained and euthanized under humane conditions in accordance with the guidelines of the Canadian Council on Animal Care. Groups of 15-week-old K18-hACE2 transgenic mice (n = 4 per group; Jackson Laboratory, 034860) were immunized with 5 ug of LNP-encapsulated or 50ug naked Gemini-D or Gemini-R formats expressing the B.1.617.2 (Delta) spike protein variant of the SARS-CoV-2 virus or the eGFP or 50ug naked Gemini-D or Gemini-R formats expressing B.1.1.529 (Omicron) Spike variant of SARS CoV-2 Spike vaccine (see Appendix Figure S1 for the sequence). Optimal doses were determined in prior experiments utilizing other Spike constructs. Mice were immunized by intramuscular injection into the right caudal thigh muscle. Blood samples were taken from the left lateral saphenous vein before vaccination at day 1 and day 28 or day 42 post-initial vaccination. During the study, mice were monitored weekly (or more frequently if needed after injections or blood collection) for any behavioural changes or changes to body condition or weight. A humane end point was determined as a 20% overall weight loss or 10% weight loss from previous weight.

### SARS-CoV-2 spike ELISA protocol

SARS-CoV-2 super stable trimer spike protein (ACROBiosystems, SPN-C52H9-50UG) was diluted to 100 ng/mL and coated onto 96 well plates using coating buffer (0.1 M Carbonate, pH 9.5). After overnight incubation at 4 °C, plates were washed four times with washing buffer (0.1% Tween-20 in 1X PBS). Subsequently, plates were blocked with blocking buffer (2% BSA, 0.1% Tween-20 in 1X PBS) overnight at 4 °C then washed five times with washing buffer. Serum samples from mice immunized with the B.1.617.2 (Delta) spike protein variant of the SARS-CoV-2 virus were serially diluted 3-fold in blocking buffer from 1:80 to 1:2160. In each well, 100 µL of serum sample dilutions were added and plates were incubated away from light at 37 °C for 1 hour. Plates were then washed four times with washing buffer before incubating with 100 µL Goat anti-mouse HRP-conjugated secondary antibody (ThermoFisher Scientific, 31430; 1:4000 dilution in blocking buffer) at 37 °C for 1 hour. Plates were finally washed with washing buffer five times before adding 100 µl/well of TMB substrate (ThermoFisher Scientific, 34028) and incubated away from light at room temperature for 20 minutes to allow for colour development. Reaction was stopped by adding 100 µl/well of stopping solution (0.16 N H_2_SO_4_). Chemiluminiscence of the plates were read using ELISA plate reader at 450 nm. A B.1.617.2 (Delta) SARS-CoV-2 spike antibody (ACROBiosystems; S1N-S58A1) was used to set up a standard curve which was structured using Graphpad prism (Version 9.4.1) from which the unknown antibody values were interpolated and the results were expressed in ng/ml (Figure 5B).

### eGFP ELISA protocol

GFP protein (Thermofisher Scientific, A42613) was diluted to 100 ng/mL and coated onto 96 well plates using coating buffer (0.1 M Carbonate, pH 9.5). After overnight incubation at 4 °C, plates were washed four times with washing buffer (0.1% Tween-20 in 1X PBS). Subsequently, plates were blocked with blocking buffer (2% BSA, 0.1% Tween-20 in 1X PBS) overnight at 4 °C then washed five times with washing buffer. Serum samples from mice immunized with the eGFP protein were serially diluted 3-fold in blocking buffer from 1:80 to 1:2160. In each well, 100 µL of serum sample dilutions were added and plates were incubated away from light at 37 °C for 1 hour. Plates were then washed four times with washing buffer before incubating with 100 µL Goat anti-mouse HRP-conjugated secondary antibody (ThermoFisher Scientific, 31430; 1:4000 dilution in blocking buffer) at 37 °C for 1 hour. Plates were finally washed with washing buffer five times before adding 100 µl/well of TMB substrate (ThermoFisher Scientific, 34028) and incubated away from light at room temperature for 20 minutes to allow for colour development. Reaction was stopped by adding 100 µl/well of stopping solution (0.16 N H_2_SO_4_). Chemiluminiscence of the plates were read using ELISA plate reader at 450 nm. A GFP antibody (Thermofisher Scientific, GFP-101AP) was used to set up a standard curve which was structured using Graphpad prism (Version 9.4.1) from which the unknown antibody values were interpolated and the results were expressed in ng/ml (Figure 5C).

### Frequency of genomic integration

The frequency of vector integration in the mouse genome was measured by a method previously described (15) (Figure 5D). Genomic DNA was extracted from the leg muscles injected with either D- or Gemini-R using tissue/cell lysis buffer (10mM Tris-Cl pH8.0, 0.1M NaCl, 10mM EDTA, 0.5% SDS), phenol/chloroform/isoamyl alcohol (Invitrogen, 25:24:1, v/v), additional chloroform extraction and ethanol/Sodium acetate precipitation. DNA was subjected to the neutral-neutral 2D gel which separates the linear mouse genomic DNA from any extrachromosomal DNA in the cells. Briefly, 10ug of genomic DNA was run in 0.4% agarose at 1 V/cm for 18 hours without ethidium bromide (EtBr), and a second dimensional electrophoresis was run in 1% agarose with EtBr at 5 V/cm for 3.5 hours. The DNA bands were then excised from the gel and purified using QiaxII gel extraction kit (Qiagen, Cat. No.20021) according to the manufacturer’s manual.

The gel extracted genomic DNA was further digested by the AscI restriction enzyme (R0558S, NEB) to make sure all the vaccine DNA was eliminated. The pure genomic DNA was separated from any extrachromosomal vaccine DNA by 2D gel electrophoresis and further digested by the AscI restriction Enzyme (R0558S, NEB) since it cuts only DNA propagated in *E. coli* cells but not genomic DNA due to the different methylation systems in prokaryotic and eukaryotic cells. As plasmid DNA lacks the origin of replication, it is not replicated by the eukaryotic cell machinery and hence it does not undergo CpG methylation. Using this approach, the remaining DNA should represent the mouse genomic DNA population. Furthermore, since the AscI site is located between NSP4 and eGFP, the remaining Gemini 1.0 DNA after successful digestion with AscI should not contribute to PCR amplification.

To determine the copy number correctly, real time q-PCR was performed using the SensiFAST™ SYBR® Kit according to the manufacturer’s instruction (BIO-94050, Bioline) and two pairs of primers (NSP4/eGFP forward primer: 5’-GTGCAAGGCAGTAGAATCAAG-3’, NSP4/EGFP reverse primer: 5’-GATGAACTTCAGGGTCAGCTT-3’ and ABCF1 forward primer: 5’-GCCGTCATCTGGCTCAATAA-3’ and ABCF1 reverse primer: 5’-CCTGCTTCTCGTACTGCTTTAG-3’. Each sample was conducted in triplicates and were PCR amplified on an Applied Biosystem 7600; the Ct values from all the samples were analyzed for the expression of eGFP and a single copy endogenous gene, ABCF1.

To calculate the integration frequency accurately, the following rationale was considered. One microgram of genomic DNA has the total of genomic DNA from 166,666 cells/0.5 genomes because the average yield from the single cell is 6pg (28). Ct values from the AscI treated samples was chosen to the calculate the copy number of Gemini 1.0 DNA spanning from NSP to eGFP in comparison to the Ct value from the standard curved created from serially diluted vaccine DNA (see Appendix Fig S1, Table S1).

### Statistical analysis

Vaccine data was first analysed for significant outliers in Graphpad prism (Version 9.4.1) using Grubbe’s test. This data was then analysed using Psych package in RStudio (R version 4.2.0). The resulting summary statistics were used to assess skewness and kurtosis of data distribution. Shapiro-Wilk and Kolmogorov-Smirnoff tests were performed in R to measure the parameters of normal distributions. Normally distributed data was subjected to *t* test (for two groups) (Figure 3D, top and bottom panels) or Analysis of Variance (ANOVA) statistical test (both using the Graphpad Prism software, version 9.4.1) along with Tukey’s and Dunnett’s post-hoc tests to test the differences between more than two groups (Figure 5B, 5C). Each point on the figure denotes individual animal in the experiment. *p*-values less than 0.05 were considered significant using 95% confidence intervals.

### Diagrams

All diagrams were created with BioRender.com.

## Acknowledgments

We would like to thank both Drs. Eliana Al Haddad and Giorgia Caspani for editorial assistance and comments on the manuscript.

## Author contributions

Conceptualization: WAJ

Vector Design: KBC, PB, WAJ

Experimental Design: KBC, JY, SK, EH, CGP, WAJ

Methodology: KBC, JY, SK, EH, SYQ, EG, IS, WAJ

Investigation: KBC, JY, SK, EH, SYQ, EG, TW, IS

Visualization: KBC, JY, SK, EH, SYQ, EG, PB, CGP, IS, WAJ

Funding acquisition: CGP, PL, WAJ

Project administration: CGP, PL, TW, WAJ

Supervision: WAJ

Writing – original draft: WAJ

Writing – review & editing: KBC, JY, SK, EH, SYQ, EG, PB, CGP, PR, IS, WAJ

## Data and materials availability

All data are available in the main text or the supplementary materials

## Supplementary Materials

Materials and Methods

Figs. S1 to S5

Table S1

