## Supplementary figs for "A Binary RNA and DNA Self-Amplifying Platform for Next Generation Vaccines and Therapeutics"

Supplementary Materials

**Appendix 1-Supplementary Figures**

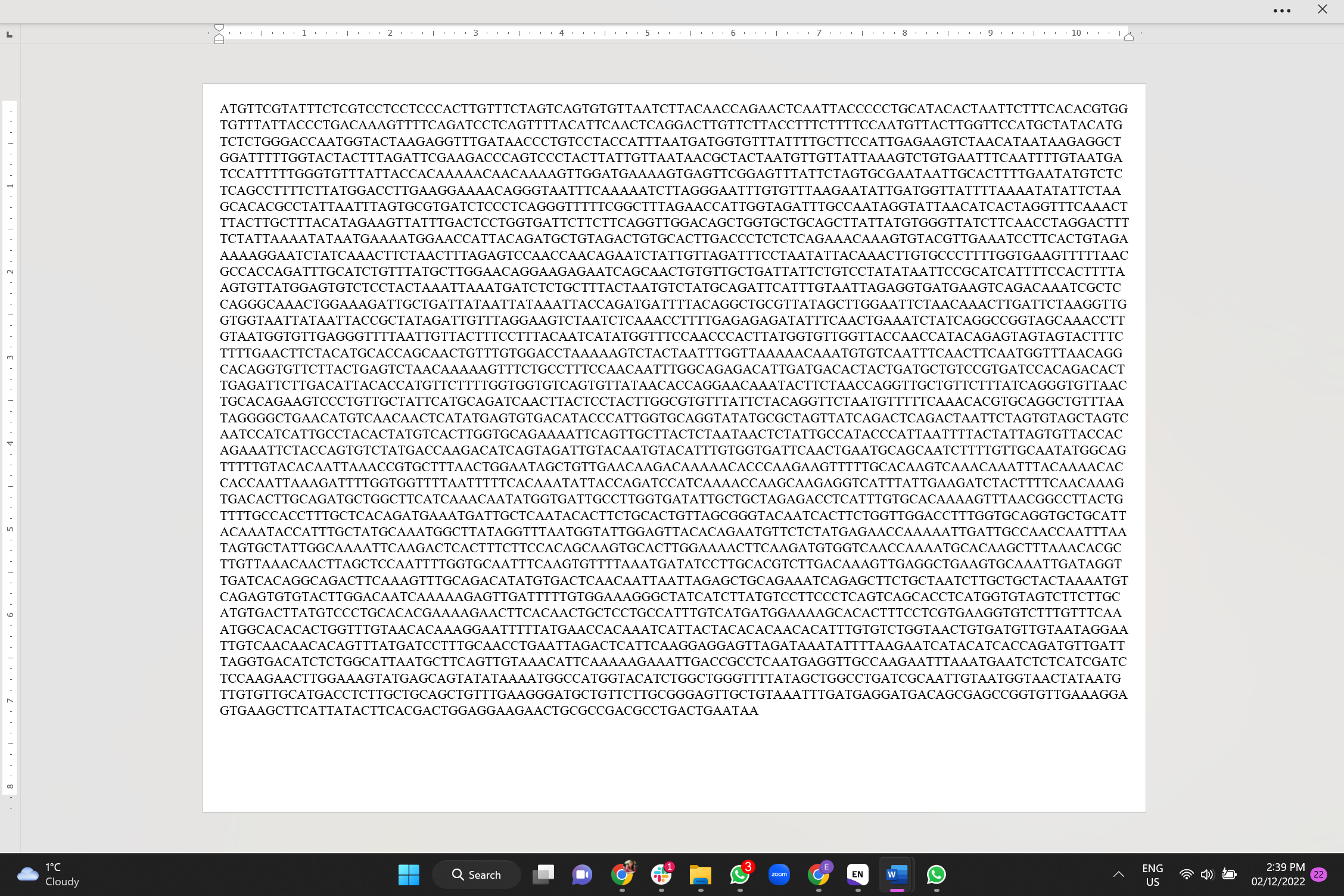

Appendix 1, Figure S1: Sequence of B.1.617.2 (Delta) spike protein variant of the SARS-CoV-2 virus

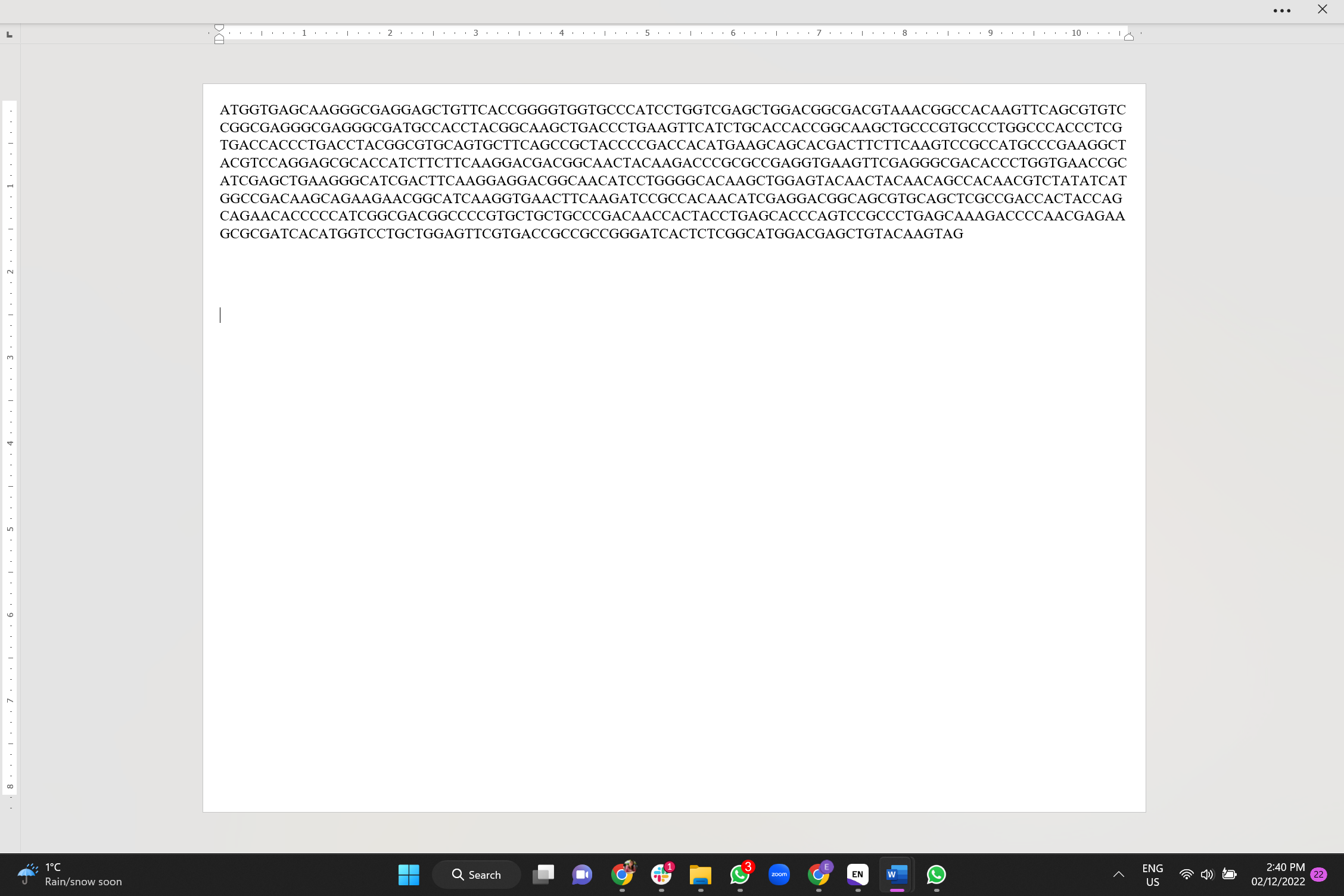

Appendix 1, Figure S2: Sequence of GEMINI-eGFP

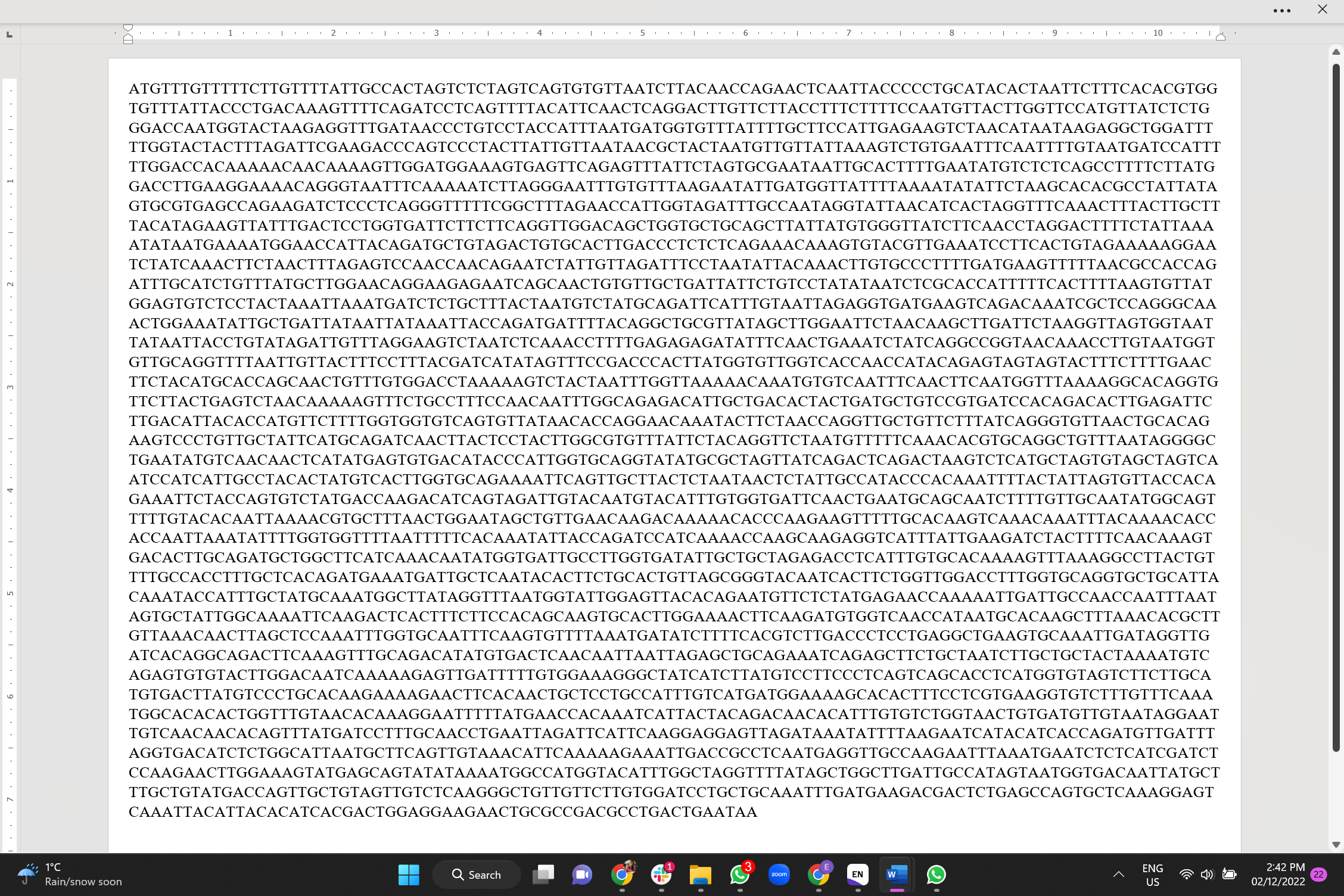

**Appendix 1, Figure S3:** **Sequence of B.1.1.529 (Omicron) Spike variant of SARS CoV-2 Spike.**

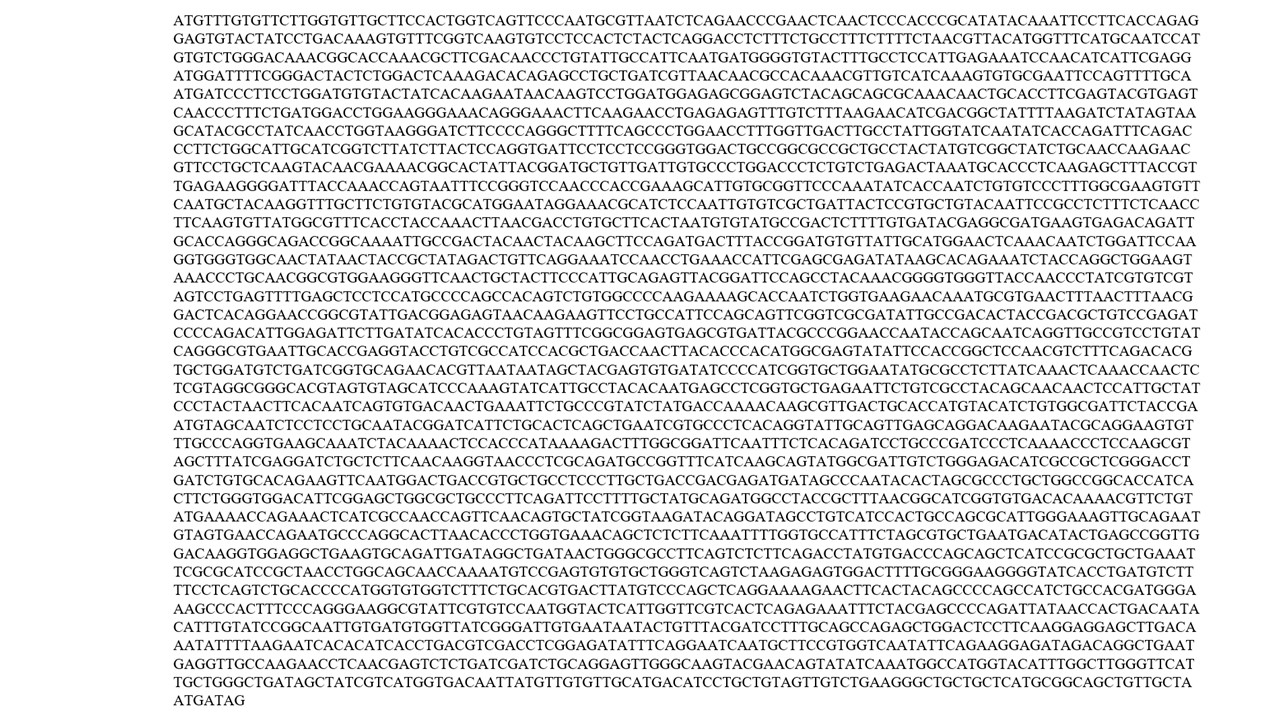

Appendix 1, Figure S4: Sequence of B.1.617.2 (Delta) spike protein variant of the SARS-CoV-2 virus from ProMab Biotechnologies, Inc.

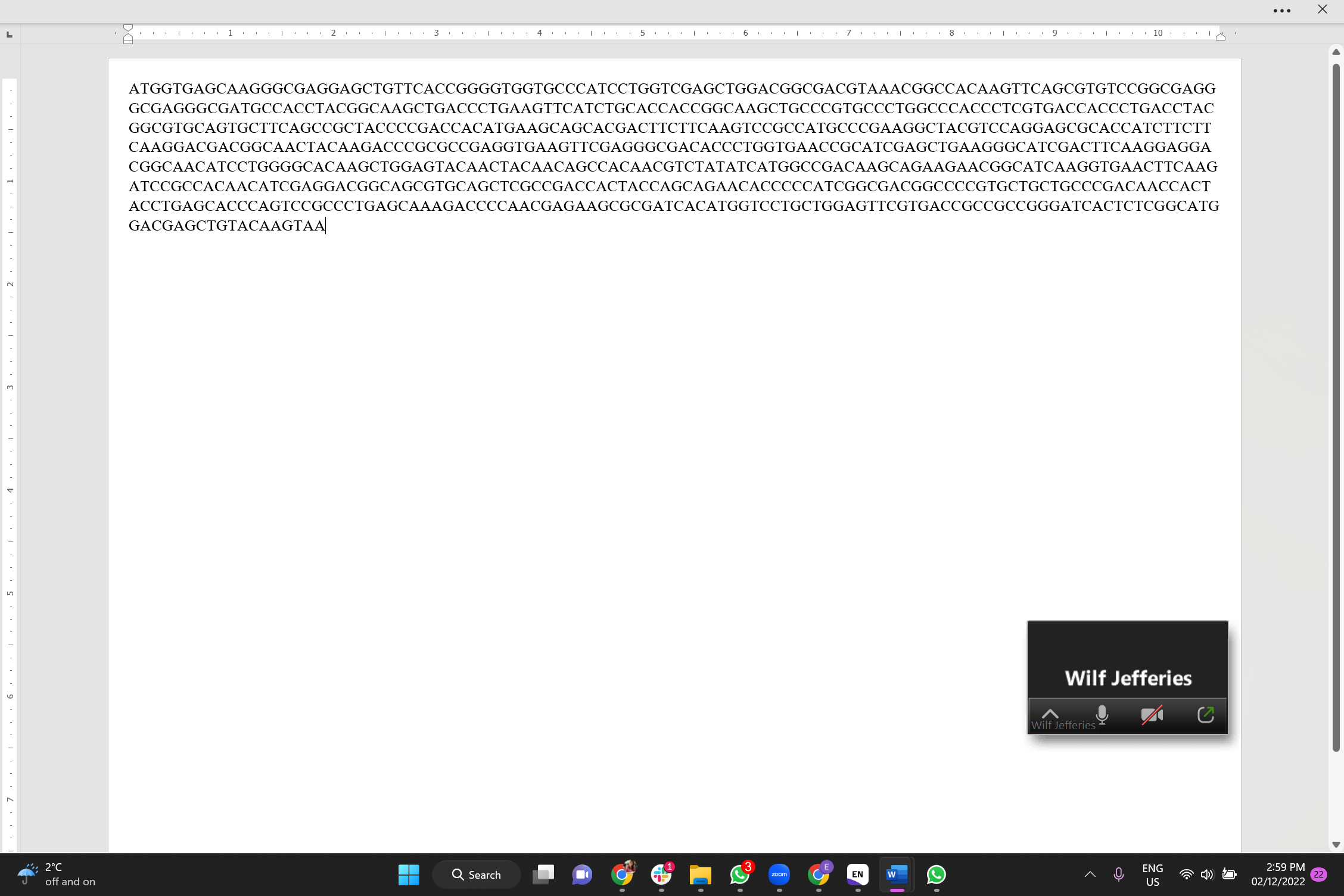

Appendix 1, Figure S5: Sequence of eGFP from ProMab Biotechnologies, Inc.

**Supplementary Tables**

Appendix 1, Table S1.

Comparison of mutation rates in different genomes and delivery systems (see Materials and Methods for a description of the protocols used to compute the figures shown for Gemini-D and Gemini-R).

| **Organism /**  **Vector** | **Spontaneous Mutation or Integration Rate** | **References** |
| --- | --- | --- |
| **Human** | 3.84 germline mutation per genome per generation  89.6 somatic mutation per genome per generation | (27, 28) |
| **Mouse (C57/B6)** | 0.945–1.46 germline mutation per genome per generation  1188 somatic mutation per genome | (27, 28) |
| **Adenovirus** | 6.7×10^-5^ integrations per genome in transduced hepatocyte | (26) |
| **Plasmid DNA vaccines (IM)** | 5×10^-5^ integrations per genome | (25) |
| **Self-amplifying DNA platform in this paper (Gemini-D)**  **Self-amplifying RNA platform in this paper (Gemini-R)** | 3×10^-6^ integrations per genome  0 integrations per genome 0 |  |
